# QTL Mapping of Seed Fe Concentration in an Interspecific RIL Population Derived from *Lens culinaris* × *Lens ervoides*

**DOI:** 10.1101/2023.06.01.543254

**Authors:** Rajib Podder, Tadesse S. Gela, Kirstin E. Bett, Albert Vandenberg

## Abstract

Biofortification of lentil (*Lens culinaris* Medik.) was investigated to potentially increase bioavailable iron (Fe) in the human diet. This study assessed the genetic variation for seed Fe concentration (SFeC) and identified the genomic regions associated with SFeC in an interspecific mapping population derived from crossing between *L. culinaris* cv. ‘Eston’ and *L. ervoides* accession IG 72815. A total of 134 RILs were evaluated in three environments. The SFeC data for individual environments and best linear unbiased prediction (BLUP) of the SFeC across environments were used for QTL analysis. The seeds of the RILs exhibited variation for SFeC from 47.0 to 102.9 mg kg^-1^ and several RILs showed transgressive segregation for SFeC. QTL analysis identified two QTLs on chromosomes 2 and 6 that accounted for 11.9-14.0% and 12.5-20.5%, respectively, of the total phenotypic variation for SFeC. The SNP markers linked to the identified QTLs may prove useful for increasing SFeC via marker-assisted selection. RILs with high SFeC can be incorporated into the lentil breeding program to broaden the genetic base of the breeding pool and/or used for the development of genetic resources for future genomic studies.

## INTRODUCTION

Iron (Fe) deficiency is one of the world’s most prevalent health concerns, especially in developing countries where diets are Fe-deficient. One fourth of the total world population is affected by anemia, an indirect indicator of Fe deficiency (McLean et al., 2009). The severity of anemia is much higher in developing countries due to an inadequate supply of nutritionally balanced foods in the context of high population growth rates, diverse food habits, and the socio-economic standing of populations. Fe is needed to regulate many metabolic processes, and since the human body cannot produce it, adequate amounts of bioavailable Fe are necessary in the diet to avoid the risk of Fe deficiency.

Among food legumes, lentil (*Lens culinaris* Medik.) is an important grain legume that provides a substantial amount of protein, complex carbohydrates, and micronutrients, including Fe (DellaValle et al., 2013). Lentil is increasingly deemed a whole food and is becoming more popular to consumers worldwide due to the presence of considerable amounts of Fe (73 –90 mg kg^-1^), Zn (44 –54 mg kg^-1^), Se (425–673 µg kg^-1^) and relatively low amounts of the micronutrient absorbance inhibitor phytic acid (2.5–4.4 mg g^-1^) (Thavarajah et al., 2011). Additional nutrients from lentil include amino acids, vitamins, phenolic compounds, dietary fiber, and resistant and slowly digestible starch, making lentil one of the healthiest foods (Tosh et al., 2013). This crop is consumed as a staple food in some developing countries where malnutrition due to Fe deficiency is more pronounced. Improving the bioavailable Fe content of traditional crops such as lentil through biofortification, a genetic approach, could be cost-effective as part of a long-term strategy to address this malnutrition.

The concept of biofortification of staple seed crops is predicated on the idea that sufficient variability for Fe concentration exists in the available gene pool of the crop. For many grain crops, research is underway to potentially increase seed iron concentration (SFeC) and bioavailability through biofortification using both genetic and agronomic methods. Limited investigation has occurred in the area of genetic strategies for increasing the bioavailability of Fe in the lentil crop through the use of genetically available gene pool of the crop.

Genes conferring regulation of Fe uptake in food crops are now identified using molecular, genetic, and biochemical techniques. Kobayashi & Nishizawa, (2012) reviewed representative genes that influence Fe deficiency in both monocot and dicot plants. These genes affect Fe uptake, translocation, subcellular translocation, and regulation in response to Fe shortage or excess at the molecular level. The lentil gene pool consists of one cultivated species, *Lens culinaris*, plus six wild species in the primary (*L. orientalis, L. odemensis, L. tomentosus),* secondary (*L. lamottei*), tertiary (*L. ervoides*) and quaternary (*L. nigricans)* gene pools (Wong et al., 2015). The wild lentil gene pool has not been extensively investigated from the standpoint of its potential contribution to nutritional improvement of cultivated lentil. The interspecific recombinant inbred line (RIL) population LR-26, developed from a cross between *L. culinaris* Eston and *L. ervoides* IG 72815 (Tullu et al., 2013), for the study of lentil disease resistance, was used in this study. The RILs were found to be very diverse phenotypically and were used to study the inheritance of many agronomically important traits (Tullu et al., 2013; Chen, 2018, Gela et al., 2021a). There is no information on whether or not the progeny and the genotypes used as interspecific parents might be of value in improving SFeC or Fe bioavailability.

Environmental factors and agronomic practices can interact with plant gene expression, which in turn can play a substantial role in differential micronutrient accumulation from soil (Bouis & Welch, 2010). Apart from genotypic variation, the lentil production environment, such as geographical location, soil quality, soil fertility, temperature, and other environmental conditions, has significant effects on micronutrient concentrations in lentil (Thavarajah et al., 2010; Kumar et al., 2013). Zhu et al. (2008) reported that the phenotype, or the observable expression of a particular complex trait such as Fe, can be influenced by numerous quantitative traits and their interactions with the environment, as well as the interaction between QTLs and the environment. Using biofortification strategies could be an attractive way to develop or explore new germplasm that can take up more Fe from the soil for deposition in seeds. It would be useful to reliably identify stable genotypes with reliably higher concentrations of bioavailable Fe. This breeding strategy would be more applicable if aided by marker-assisted selection (MAS). However, limited research has been done to uncover the genomic regions (QTLs) harbouring genes for high Fe uptake in lentil seeds or to identify closely linked molecular markers to screen and select the most favorable genotypes. Aldemir et al. (2017) mapped QTLs conferring SFeC on linkage groups (LG) 1, 2, 4, 5, 6, and 7 using the GBS approach in a cultivated lentil RIL population. Similarly, previous association mapping studies (Khazaei et al., 2017; Singh et al., 2017; Kumar et al., 2019) identified molecular markers that were associated with SFeC. QTL/genes conferring Fe uptake in food crops are mostly identified in cereal crops (Stangoulis et al., 2007, Xu et al., 2012; Jin et al., 2013; Srinivasa et al., 2014). Kobayashi & Nishizawa, (2012) reviewed representative genes that influence Fe deficiency in both monocot and dicot plants. These genes affect Fe uptake, translocation, subcellular translocation, and regulation in response to Fe shortage or excess at the molecular level.

In comparison to cultivated lentil, *Lens ervoides* has proven to be a good source of resistance genes for various lentil diseases (Tullu et al., 2010; Vail et al., 2012; Podder et al., 2013; Gela et al., 2020, 2021b, Gela, 2021, Adobor et al., 2022). It is important to ascertain how SFeC may vary in these interspecific lentil hybrids and their progenies, which can contribute genetic diversity to cultivated lentil breeding. The fundamental question is whether or not interspecific hybridization results in the development of lentil germplasm with more variation in SFeC, which would be essential to make progress in biofortification. This type of information has never been reported. The present study was conducted to assess the genetic variation of SFeC and identify the genomic region(s) associated with SFeC in an interspecific RIL population of lentil. To our knowledge, this is the first study that investigates the inheritance and genomic regions conferring SFeC in interspecific hybrids of lentil.

## MATERIALS AND METHODS

### Plant material

A mapping population of 134 interspecific RILs from the LR-26 population developed from a cross between *L.culinaris* cv. ‘Eston’ and *L. ervoides* accession IG 72815 (Tullu et al., 2013) was used to evaluate SFeC. The RILs were developed from F_2_ by single seed descent, and the F_7_- derived bulked seeds of the RILs were selfed for at least three additional generations (Tullu et al., 2013; Gela et al., 2021a) and have been maintained at the Crop Development Centre (CDC), University of Saskatchewan (USask).

### Field experiments

The RIL population and parents were evaluated at the Sutherland farm (STH; 52°15′ N, 106°52′ W) in 2015 and at the Crop Science Field Lab (CSFL; 52°36′ N, 106°62′ W) in 2014 and 2015. The experiments were designed as hill plots arranged in a randomized complete block design (RCBD) with three replications in all three environments. A total of 20 seeds of each RIL genotype, including the parents, were sown at depth of ∼3.8 cm using a 4-row tray hill plot planter. Each hill plot in a row had a different genotype, with a plot length of 30 cm and a spacing of 30 cm between rows. To increase field germination, all seeds of each genotype, except for Eston, were scarified and stored at -20°C for two days before seeding. Rainfall, maximum, and minimum temperature data were collected from Environment Canada (https://climate.weather.gc.ca/index_e.html). Soil samples were collected from different parts of each experimental site for both locations and analysed by ALS Laboratory Group Agriculture Services, Saskatoon, Canada (**Supplementary Tables 1 and 2**).

### Seed harvest and Fe analysis

The plant growth habit of all lentil species is indeterminate (Knowpulse, 2021), and unlike cultivated species, wild accessions have a dehiscent pod trait that causes seed dispersal at pod maturity. Collection of seeds, therefore, requires extra care and techniques that minimize seed loss (Figure 1). Every plant in each hill was enclosed with a mesh bag prior to harvest. The lower end of the mesh bag was tied at the bottom of the plant so that shattered seeds accumulated inside the bag. The top portion of the mesh bag was kept open and tied with nylon rope to hold the mesh bag in an upright position and to provide adequate sunlight and aeration (Figure 1). The harvested seeds were stored at room temperature prior to the estimation of SFeC. The Fe concentration in the lentil seed samples was analyzed by flame atomic absorption spectrophotometry (F-AAS, Nova 300, Analytic Jena AG, Konrad-Zuse-Strasse, Neu-Ulm, Germany) as described in a previous study (Podder et al., 2017; Diapari et al., 2014). In brief, 0.5 g of each seed sample was digested in a 30-mL digestion tube with HNO_3_-H_2_O_2_ using an automatic digester (Vulcan 84, Questron Technology, Ontario, CA, USA). A total of 72 samples were digested in each run, including eight standards (yellow lentil laboratory checks) and four blanks. Samples were first digested with HNO_3_ at 90°C for 45 min, followed by the addition of 5 mL of 30% H_2_O_2_, and then further digested for another 65 min. The solutions were then reduced with 3 mL of 6 M HCl, followed by heating at 90°C for 5 min prior to cooling to room temperature. All sample solutions were then diluted with deionized water to a volume of 25 mL. Six mL of each of the digested samples was then used to determine the SFeC. The digestion was repeated twice, with three technical replications per repeat. The mean for the technical replicates was calculated and used for further statistical analysis.

**Figure 1.**
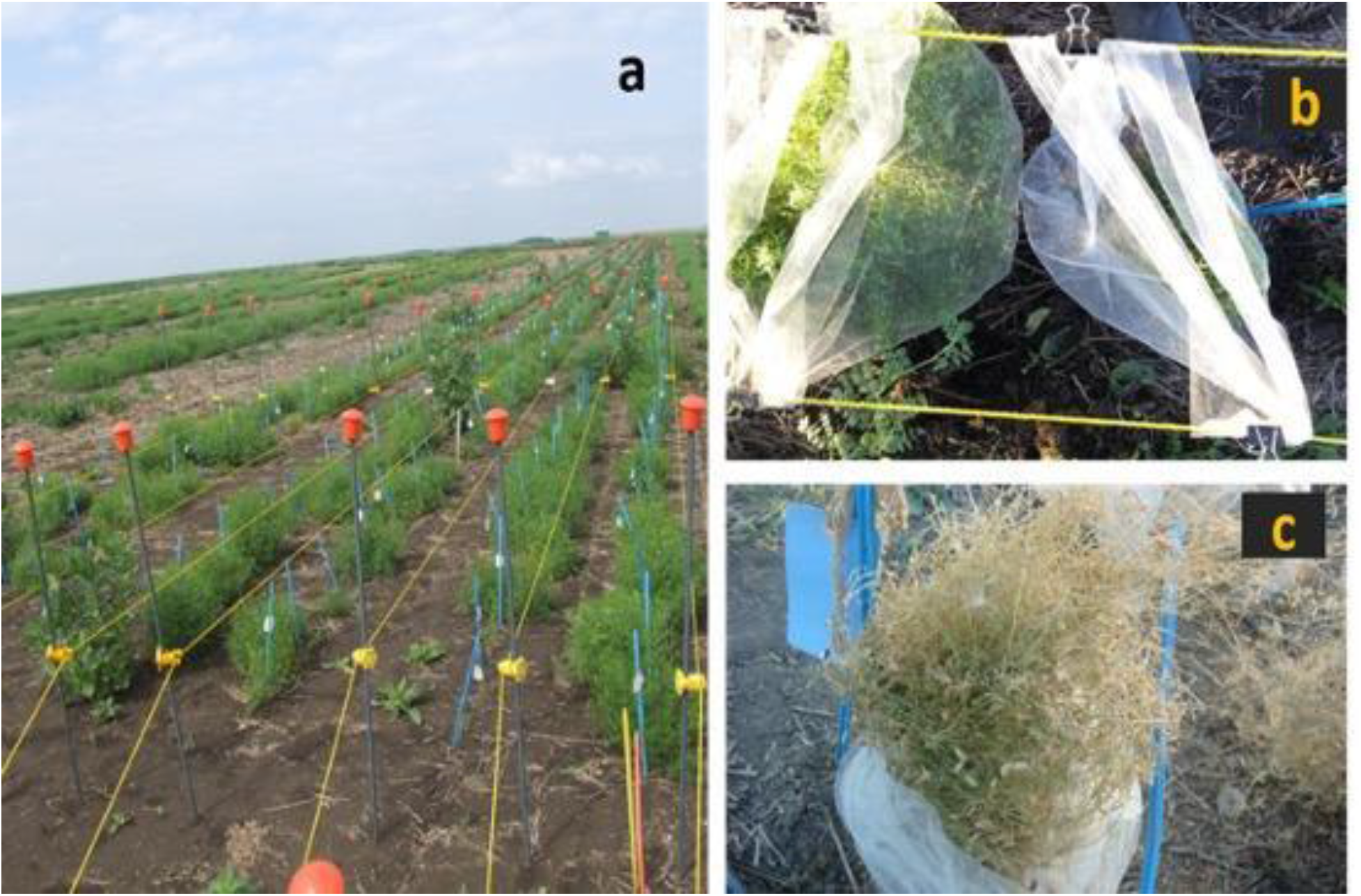
Images showing (a) field view of plants at mid-season growth stage in hill plots, (b) plants inside mesh bags used for seed collection, and (c) seed harvesting techniques for wild species genotypes used for the Fe accumulation study.

### Statistical analysis

The analysis of variance (ANOVA) was performed using PROC MIXED of SAS version 9.4 (SAS Institute Inc., Cary, NC, USA). Levene’s test of homogeneity of variance (test) was performed by environments before pooling SFeC data across environments. In the combined ANOVA, genotype, environment, and the interaction of genotype × environment were treated as fixed factors, and replications nested within each environment were considered random. The significance of variances was declared at the 5% level, and least square means were estimated for genotypes using LSMEANS statements. The variance components were estimated using the PROC VARCOMP procedure of SAS 9.4. Estimates of broad sense heritability (h_B_ ^2^), both within and across environments, were calculated using the formula h_B_ ^2^ = σ^2^g / (σ^2^g + σ^2^e/r) and h_B_ ^2^ = σ^2^g / (σ^2^g +σ^2^ge /e + σ^2^e/er), respectively (Singh et al., 1993; Ubayasena et al., 2010). Where σ^2^g, σ^2^ge, σ^2^e, r and e were estimates of genotypic variance, genotype × environment interaction, residual or error variance, number of replications, and number of environments, respectively. The best linear unbiased prediction (BLUP) was estimated across environments to reduce environmental variation using SAS version 9.4.

### QTL analysis

The Fe data for the three environments individually, and the BLUP data for the combined analysis were used for detecting the QTL associated with the SFeC. A genetic linkage map of the LR-26 population was used for QTL mapping (Gela et al., 2021a). The map consisted of 5455 SNP markers and spanned a total genetic map distance of 3252.8 cM at an average marker interval of 0.6 cM. Genotyping information for the LR-26 genetic linkage map can be found through the KnowPulse database, accessible at: http://knowpulse.usask.ca/Geneticmap/2691115 (accessed 2^nd^ March 2023). The QTL analysis was performed using the inclusive composite interval mapping (ICIM) implemented in QTL IciMapping v.4.2 software (Meng et al., 2015). The genome was scanned with a walking step of 1 cM along the chromosome. The significant threshold logarithm of the odds (LOD) scores for detection of the QTL were calculated based on 1000 permutations at P ≤ 0.05. Mapchart (Voorrips, 2002) was used for the graphical representation of the identified QTLs on the genetic linkage map.

## RESULTS

### Soil Fe status and weather conditions

The climate data for both years were collected from the Environment Canada (2017) website. The average monthly temperature (°C) was similar across the three environments from May to August (Supplementary Table 1). Total precipitation (mm) was 65.4 mm higher in 2014 (wetter than average) compared to 2015. The characteristics of soil samples collected from different parts of each experimental site are shown in Supplementary Table 2.

### Seed Fe concentration

A wide range of variation was observed for SFeC among the lines in the LR-26 RIL population (Figure 2; Supplementary Table 3). The range in SFeC among the RILs was 47.0 to 102.9 mg kg^-1^. The genotype, environment, and genotype × environment effects were highly significant (p ≤ 0.001) across the environments (Table 1). The mean SFeC (mg kg^-1^) per environment for LR-26 RILs varied from 67.0 to 76.4, with an overall mean of 71.3 SFeC (mg kg^-1^). The mean SFeC for the Sutherland (STH 2015) was higher than the CSFL 2014 and 2015. Higher broad-sense heritability estimates for SFeC across the environments were observed, ranging from 0.85 to 0.94 and 0.87 for combined data. A significant (p ≤ 0.05) negative correlation was observed for SFeC and seed yield (Supplementary Figure 1). Although the Fe accumulation of the two parents is comparable, the *L. culinaris* parent Eston had a higher SFeC (65.1 mg kg^-1^) than the *L. ervoides* parent accession IG 72815 (57.7 mg kg^-1^) (p ≤ 0.05). Among lines of the LR-26 population, 13 (8%) and 82 (61%) RILs had significantly lower and higher SFeC (mg kg^-1^) than the *L. culinaris* parent Eston, respectively (Supplementary Table 3). These lines can be considered transgressive segregants. The analysis based on Pearson correlation (r) showed that SFeC were positively correlated across the three environments, with r ranging from 0.59 to 0.79 (p < 0.001).

**Figure 2.**
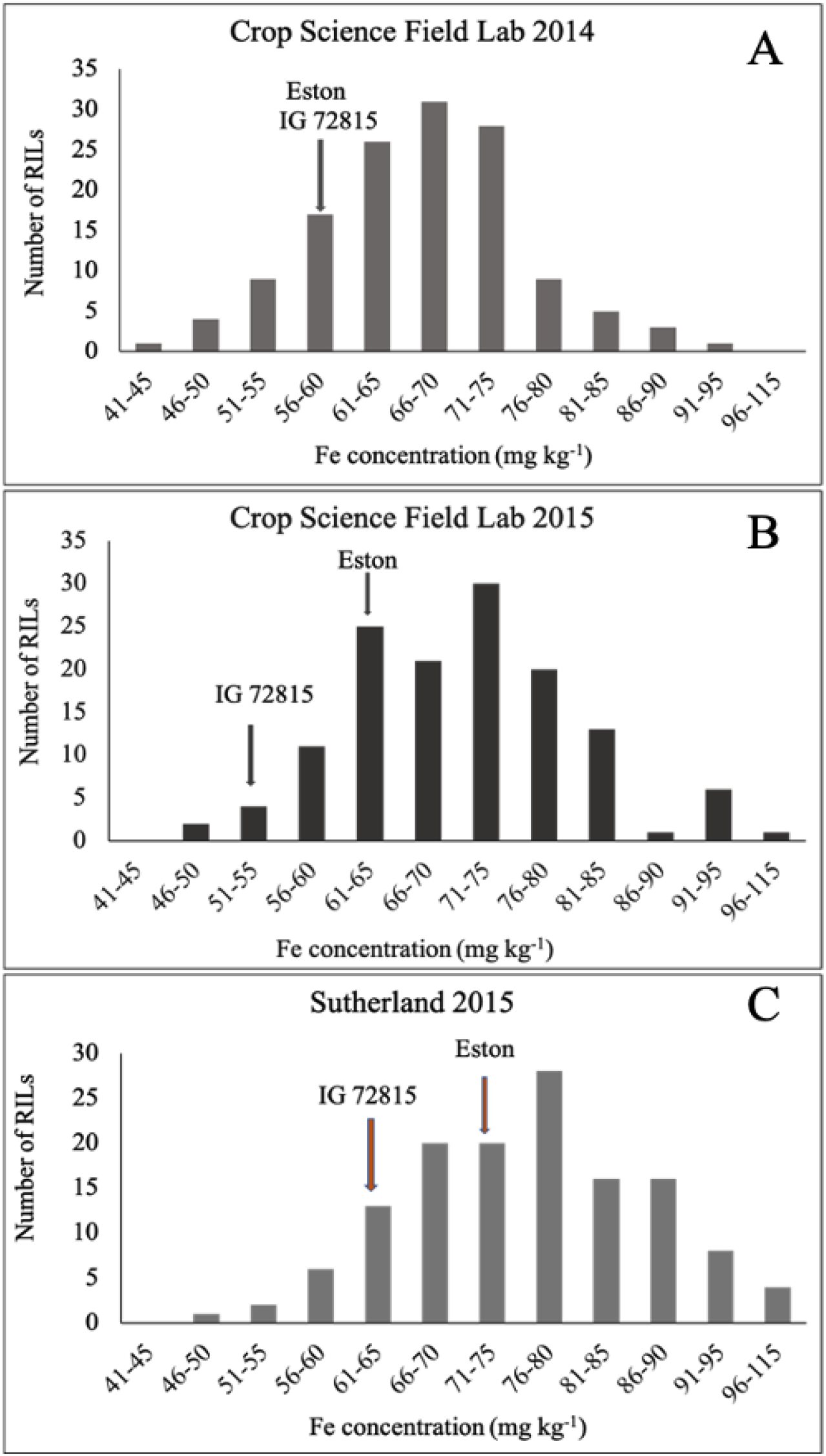
Frequency distribution of seed Fe concentration of parents and 134 interspecific lentil recombinant inbred lines (RILs) derived from Eston × IG 72815 (LR-26) grown at Crop Science Field Lab (CSFL), University of Saskatchewan in 2014 (A), Crop Science Field Lab (CSFL), University of Saskatchewan in 2014 (B), and Sutherland crop research field in 2015 (C). Seed Fe concentration of RIL parents is indicated by arrows.

**Table 1.** Analysis of variance, variance components and broad-sense heritability for seed Fe concentration (mg kg^-1^) for 134 lentil RILs (including parents) of LR-26 evaluated at Sutherland farms in 2015 and Crop Science Field Laboratory, University of Saskatchewan in 2014 and 2015.

| Environment | Mean<sup>a</sup> | Range | CV | $h_B^2$ | $\sigma^2_g$ | $\sigma^2_{ge}$ | $\sigma^2_e$ |
| --- | --- | --- | --- | --- | --- | --- | --- |
| CSFL 2014 | 66.95 | 39.42 - 100.66 | 4.61 | 0.94 | 75.15** | - | 14.3 |
| CSFL 2015 | 70.92 | 44.08 - 112.19 | 7.63 | 0.88 | 84.57** | - | 33.96 |
| Sutherland 2015 | 76.38 | 39.80 - 137.75 | 8.18 | 0.85 | 99.33** | - | 52.25 |
| Combined | 71.30 | 39.42 - 137.75 | 6.32 | 0.87 | 67.24** | 18.38** | 33.12 |

<sup>a</sup>Mean Fe concentration ( $\text{mg Kg}^{-1}$ ); \*\*\* $p < 0.001$ ; $\sigma^2_g$ , genotypic variance; $\sigma^2_{ge}$ , Genotype ×
Environment variance; $\sigma^2_e$ , error variance; $h_B^2$ , heritability; CV, Coefficient of variation.

### QTL analysis for seed Fe concentration

The mean data for the individual environments and the BLUP values across environments were used for mapping of the QTL associated with SFeC. Two QTLs were identified in the LR-26 population (Figure 3) on chromosome 2 and 6. The QTL detected at individual environments and the details of each QTL identified are presented in Table 2. QTL on a chromosome were considered the same QTL when their critical intervals overlapped or the maximum distance between the QTLs were less than 5 cM. The QTL on chromosome 2 was consistently detected in individual environments as well as with BLUP data, accounting for 11.9 to 14% of the phenotypic variation. The QTL on chromosome 6 were identified with the data from CSFL 2015, Sutherland 2015 and the BLUP data, explained 12.5 to 20.5% of the phenotypic variation. The allele for the QTL on chromosome 6 was contributed by the cultivated lentil parent, Eston, whereas the alleles for the QTL on chromosome 2 were contributed by the *L. ervoides* parent, accession IG 72815.

**Figure 3.**
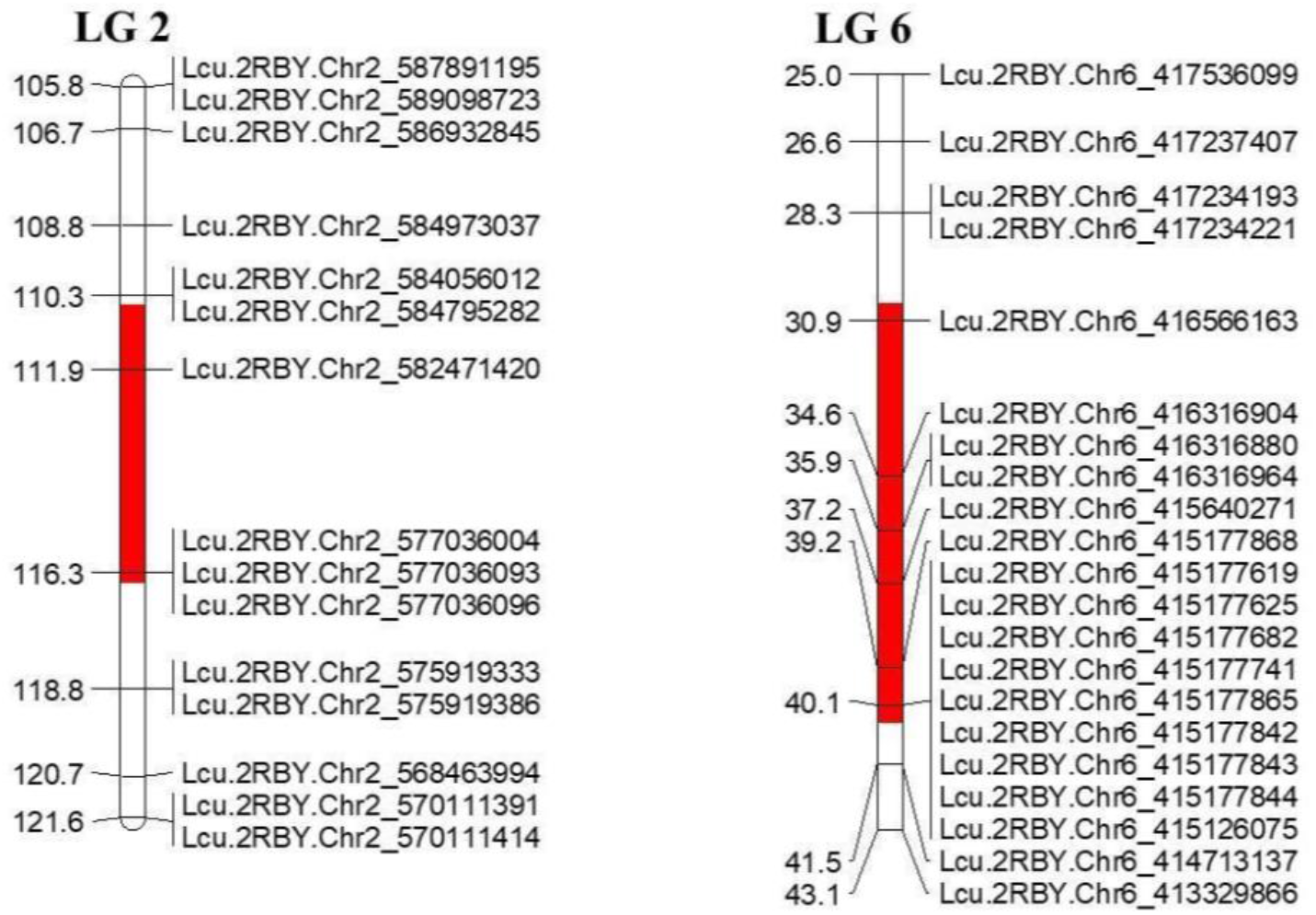
Quantitative Trait Loci (QTLs) associated with seed Fe concentration in LR-26 populations on linkage groups 2 and 6.

**Table 2.** Quantitative trait loci (QTL) for seed Fe concentration detected by ICIM-ADD analysis in the LR-26 RIL population derived from a cross between *L. culinaris* Eston and *L. ervoides* accession IG 72815.

| Environment<sup>a</sup> | QTL<sup>b</sup> | LG<sup>c</sup> | Pos<sup>d</sup><br>cM | Left marker | Right marker | LOD<sup>e</sup> | PVE<br>(%)<sup>f</sup> | Add<sup>g</sup> | Left<br>CI<sup>h</sup> | Right<br>CI<sup>h</sup> |
| --- | --- | --- | --- | --- | --- | --- | --- | --- | --- | --- |
| CSFL 2014 | qFe2 | 2 | 112 | Lcu.2RBY.Chr2_582471420 | Lcu.2BY.Chr2_577036004 | 4.43 | 13.8 | -3.1 | 110.5 | 116.5 |
| CSFL 2015 | qFe6 | 6 | 34 | Lcu.2RBY.Chr6_416566163 | Lcu.2RBY.Chr6_416316904 | 4.3 | 14.0 | 3.8 | 30.5 | 38.5 |
| STH 2015 | qFe2 | 2 | 112 | Lcu.2RBY.Chr2_582471420 | Lcu.2RBY.Chr2_577036004 | 3.5 | 11.9 | -3.7 | 110.5 | 116.5 |
|  | qFe6 | 6 | 40 | Lcu.2RBY.Chr6_415177868 | Lcu.2RBY.Chr6_415177619 | 3.6 | 12.5 | 3.7 | 39.5 | 40.5 |
| BLUP | qFe2 | 2 | 112 | Lcu.2RBY.Chr2_582471420 | Lcu.2RBY.Chr2_577036004 | 4.8 | 13.9 | -2.9 | 110.5 | 116.5 |
|  | qFe6 | 6 | 40 | Lcu.2RBY.Chr6_415177868 | Lcu.2RBY.Chr6_415177619 | 6.7 | 20.5 | 3.4 | 39.5 | 40.5 |

<sup>a</sup>Environment – CSFL (Crop science field lab), STH (Sutherland 2015), BLUP (best linear unbiased prediction), <sup>b</sup>QTL nomenclature – q=QTL, Fe = Fe concentration, 2=LG; <sup>c</sup>LG - Linkage group (chromosome); <sup>d</sup>Pos – Position of QTLs; <sup>e</sup>LOD – Logarithm of odd for each QTL; <sup>f</sup>PVE - Phenotypic variation explained; <sup>g</sup>Add - additive effect; <sup>h</sup>CI – Left and right critical interval.

## DISCUSSION

Lentil is one of the oldest cultivated crops, and its global per capita consumption is increasing much faster than other pulse crops. In some developing countries lentil is considered a partially staple food due to its nutritive value as a relatively inexpensive protein source compared to animal sources. Lentil is cultivated in many different agro-ecological regions around the world, therefore, geographical location, soil factors, temperature, and other conditions can have a significant influence on lentil SFeC (Thavarajah et al., 2010). Assessment of genotype × environment interaction for micronutrient dense germplasm is essential for determining the influence of growing environments on micronutrient content expression (Bouis & Saltzman, 2017). The interaction can also reduce the genotypic stability of micronutrient-dense genotypes. Results from this study determined the SFeC of *L. culinaris* × *L. ervoides* interspecific hybrid RILs and their parents across three environments. Significant effects of genotype, environment, and genotype × environment interactions were observed for SFeC. Gregorio, (2002) reported that environment had a significant influence on common bean (*Phaseolus vulgaris* L.) SFeC, but that high-Fe bean genotypes accumulated more Fe compared to low-Fe genotypes when grown at the same location in the same growing season. Genotype × environment interaction also increases with the increase of number of environments and the number of genotypes (Baye et al., 2011).

A continuous frequency distribution in the LR-26 RIL population indicated quantitative the inheritance of Fe uptake, transport or storage in seeds. Evidence for multiple genes related to inheritance of SFeC was reported in some studies (Upadhyaya et al., 2016; Diapari et al., 2014; Wang et al., 2008). Mineral accumulation in higher plants or in different plant parts is controlled by many genes with major or minor effects. For instance, a study of an interspecific cross between wild and cultivated species of common bean showed quantitative inheritance of SFeC (Guzmán-Maldonado et al., 2003).

In lentil, a wide range of phenotypic and genotypic variability among different lentil species was reported for above- and below-ground traits (Gorim & Vandenberg, 2017a). Particularly for root system-related traits such as total root length, total root surface area, root length per unit volume of soil, total root volume, the mean root diameter, root volume, and fine root distribution in different soil horizons (Gorim & Vandenberg, 2017b). These variations in phenotypes between the cultivated and *L. ervoides* genotypes could contribute to the quantitative distribution of the SFeC in the LR-26 RIL population. Moreover, chromosomal rearrangements after hybridization between two species can also influence the phenotypic expression of the trait (Baack & Rieseberg, 2007). The soil pH and existing Fe status, on the other hand, could have influenced Fe accumulation. There was a substantial difference in precipitation and temperature, as well as soil pH and DTPA-extractable [Fe] (mg kg^-1^) between years and locations (**Supplementary Tables 1 and 2**). A previous study reported that at naturally alkaline pH conditions, soil Fe precipitates and thus decreases its availability (Pandian et al., 2011). Thus, these non-allelic interactions may influence quantitative traits and their phenotypic expression (Saha et al., 2013).

In the present study, more than 80% of the interspecific RILs had higher SFeC compared to the *L. culinaris* parent Eston and displayed transgressive segregation. DeVecente and Tanksley (1993) emphasised that interspecific transgressive segregants are more likely to be observed when parental genotypes are more similar in trait phenotypes. Likewise, we observed a small difference in SFeC between the two parents, suggesting that both parents carried a few different genes with alleles contributing to the trait (Xu et al., 2012). Transgression could also occur as a result of complementary gene actions (Reiseberg et al., 2000), indicative of the two QTLs identified that are inherited from both parents of the LR-26 population. Some LR-26 RILs had higher SFeC but a lower seed yield. Karaköy et al. (2012) reported an inverse relationship between SFeC and both seed size and seed weight of lentil landraces.

In this study, we mapped two QTLs associated with Fe concentration in seeds of the LR-26 population. The QTLs were identified on chromosomes 2 and 6, accounting for 11.9 to 20.5% of the variance in SFeC. The QTLs were consistently detected across test environments (except CSFL 2014), supporting the high heritability estimates and the observed positive correlation between environments. Khazaei et al. (2017) reported two significantly associated SNPs with SFeC on lentil chromosome 5 and other potential genomic regions, including chromosome 6, using association mapping (AM). Similarly, using the AM approach two SSR markers (Kumar et al., 2019) and three SSR markers (Singh et al., 2017) associated with SFeC were reported. Aldemire et al. (2017) found that 21 QTL regions on six linkage groups (LG1, 2, 4, 5, 6, and 7) explained 5.9 to 14% of the phenotypic variation in SFeC using GBS-based SNP markers. Also, Blair et al. (2009) reported 13 QTLs for SFeC after studying an inter-gene pool common bean RIL population. However, other previous studies identified fewer QTLs linked to grain Fe concentration in cereal crops such as rice (Stangoulis et al., 2007), wheat (Xu et al., 2012; Srinivasa et al., 2014), and maize (Jin et al., 2013). similar to the current study.

Biofortification of lentil cultivars through plant breeding is considered a sustainable and cost-effective solution for overcoming Fe malnutrition (White and Broadley, 2005). Mapping of genomic regions responsible for high Fe content in lentil is the first step toward the development and application of marker assisted breeding. In major cereal crops, significant knowledge has been gained on the molecular mechanisms underlying the uptake of Fe (Bauer et al., 2004; Cakmak, 2002). Therefore, increasing the SFeC of lentil cultivars through breeding without affecting the yield and quality could be possibly achieved via MAS and can complement the ongoing efforts to fortify lentils (Podder et al., 2017).

## CONCLUSIONS

The objective of the current study was to evaluate the accumulation of SFeC in an interspecific RIL population (LR-26) across three environments and to identify QTLs associated with SFeC. A wide range of variation for SFeC was observed among the RILs. The RILs with high SFeC can be used in breeding programs to broaden the breeding gene pool of lentil and develop targeted populations for future genetic studies. Two QTLs were mapped on chromosomes 2 and 6, which are effective across tested environments and explain the higher broad-sense heritability estimated in individuals and across environments. The SNP markers linked to the identified QTLs will be useful to improve seed Fe content via MAS.

## AUTHOR CONTRIBUTIONS

Rajib Podder designed the study, analyzed data, interpreted results and wrote the manuscript. Tadesse Gela helped with data analysis, interpretation of results and writing the manuscript. Kirstin Bett and Albert Vandenberg contributed to critically revising the paper.

## ACKNOWLEDGMENTS

The authors are grateful for technical assistance provided by Barry Goetz, Crop Development Centre, University of Saskatchewan. The authors would like to acknowledge financial assistance received from The Saskatchewan Ministry of Agriculture (Agriculture Development Fund), Grand Challenges Canada, Saskatchewan Pulse Growers, and the NSERC Industrial Research Chair Program for their generous funding support.

---

**Supplementary Figure 1.**
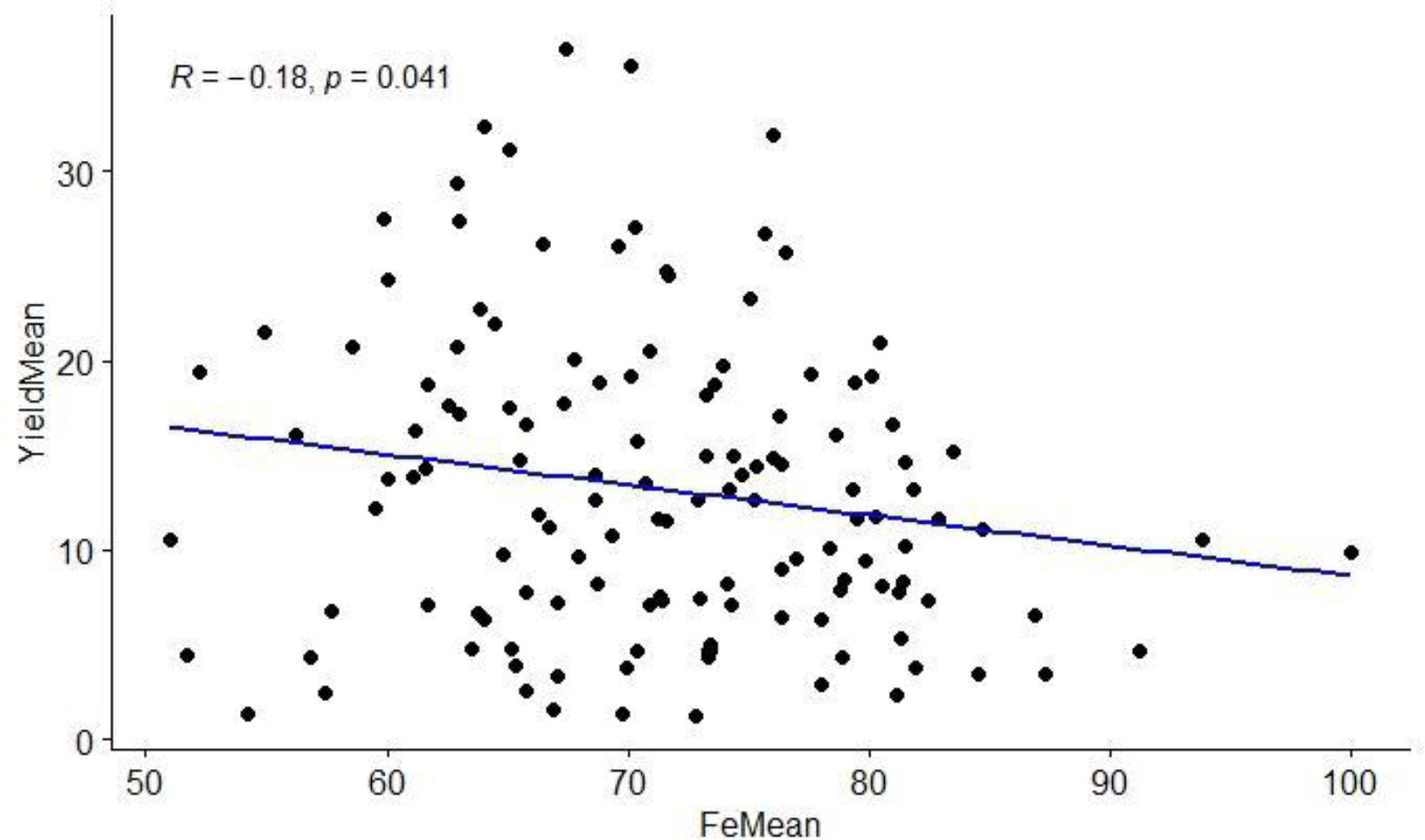
Correlation coefficient between mean seed yield (g hill-plot^-1^) and Fe concentration (mg kg^-1^) of LR-26 population evaluated at Sutherland farms in 2015 and Crop Science Field Laboratory, University of Saskatchewan in 2014 and 2015.

**Supplementary Table 1.**
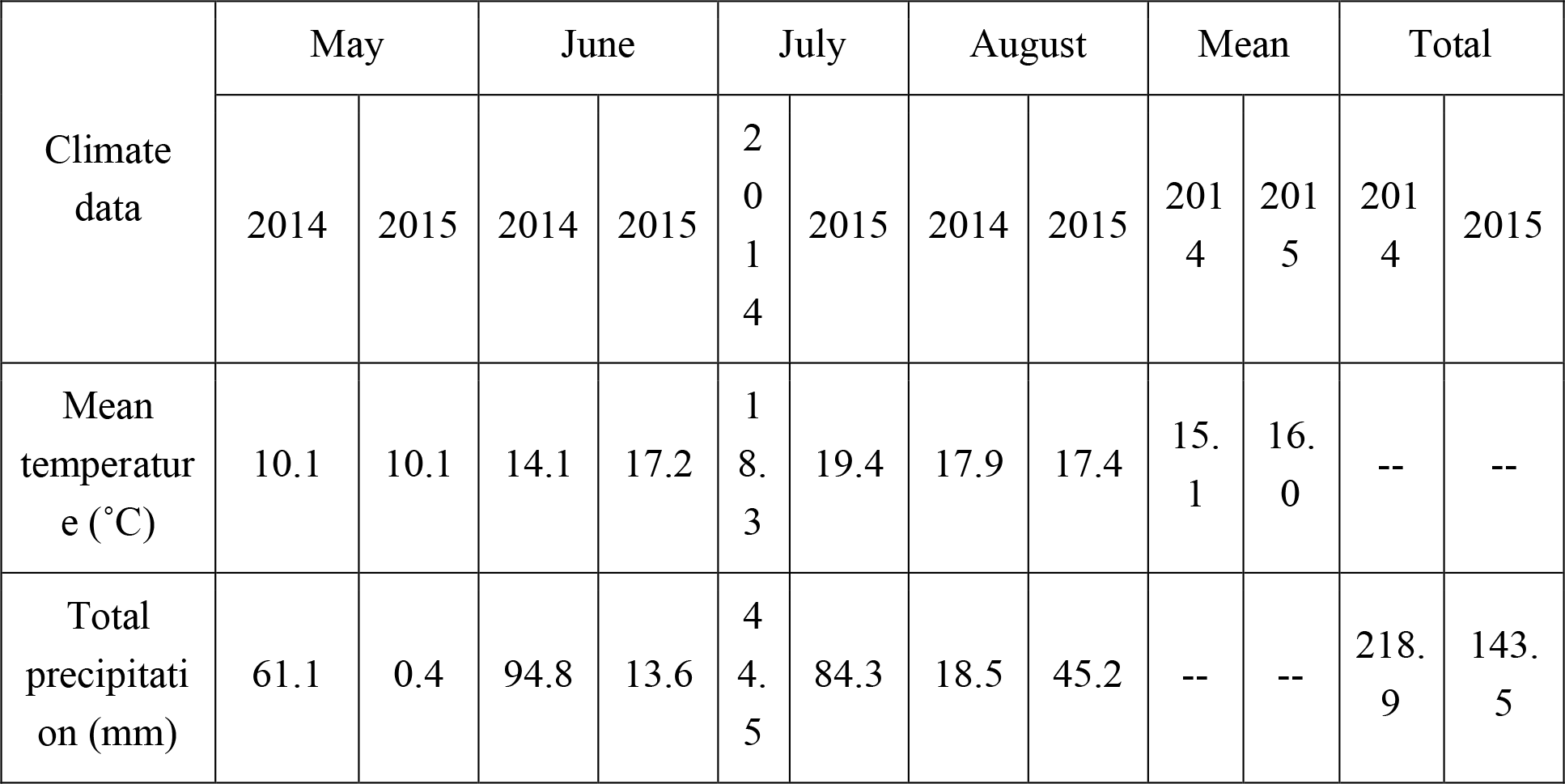
Mean temperature (°C) and total precipitation (mm) for the year 2014 and 2015 growing seasons (May-August) at Saskatoon area.

**Supplementary Table 2.** Soil analysis from the field locations at Sutherland (STH) and Crop Science Field Lab (CSFL).

| Location | Depth<br>(inches) | Texture | pH | Salinity<br>rating | Organic<br>matter (%) | DTPA-extractable<br>[Fe] (mg kg<sup>-1</sup>) |
| --- | --- | --- | --- | --- | --- | --- |
| STH | 0-6 | Clay Loam | 6.6 | NS | 3.6 | 58.5 |
| CSFL | 0-6 | Loam | 7.5 | NS | 3.7 | 45.1 |

NS, Non-saline

**Supplementary Table 3.**
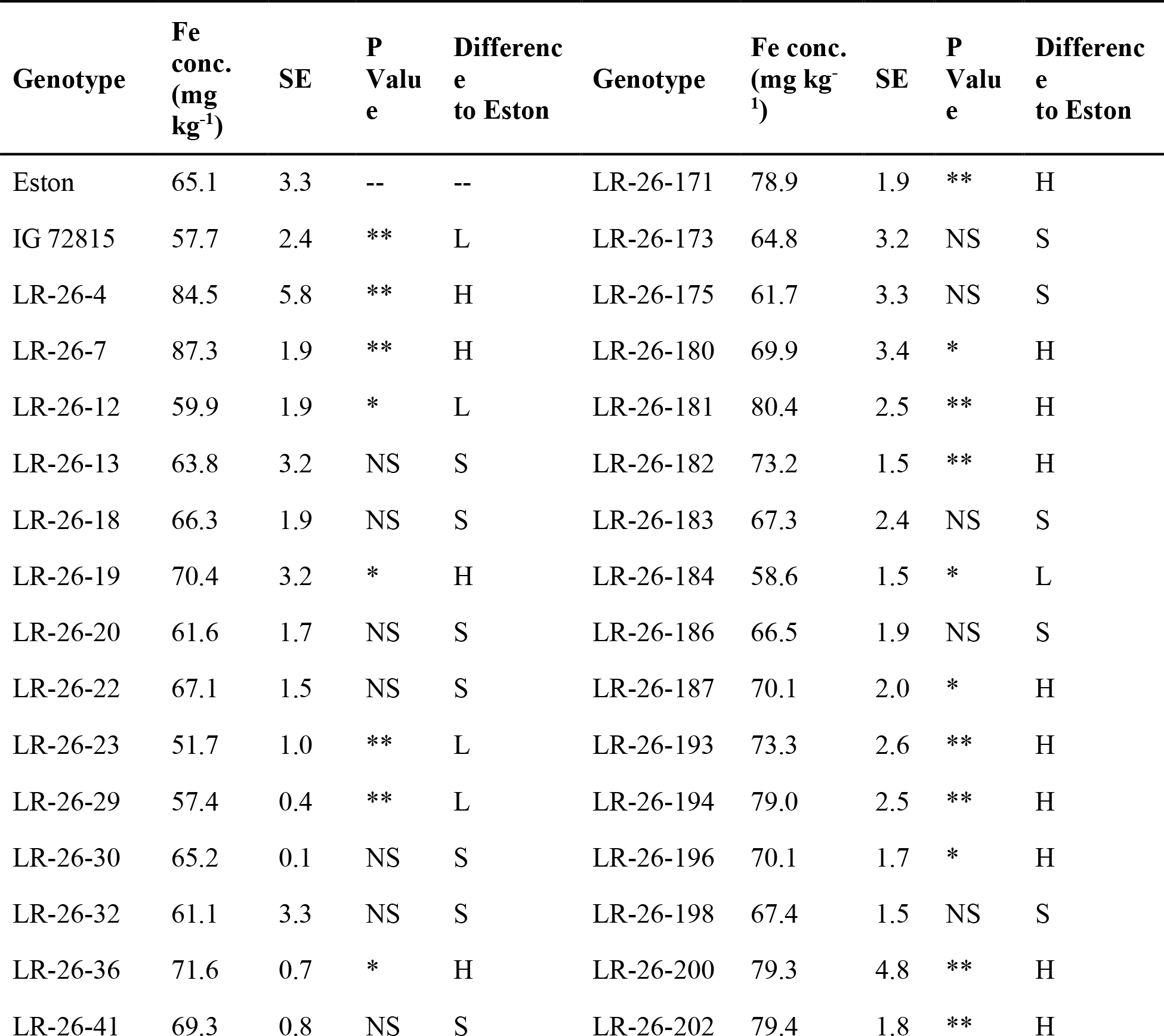
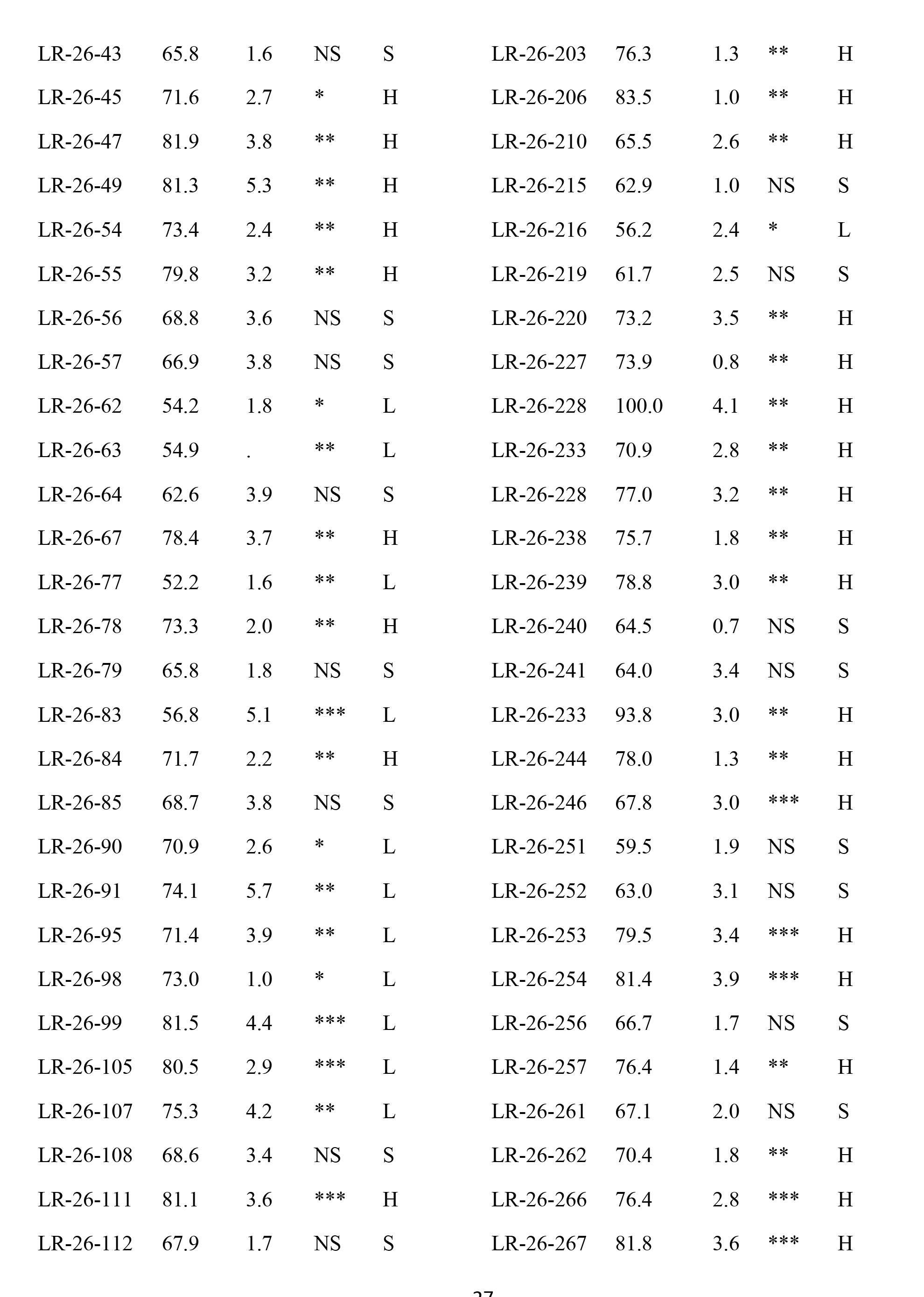
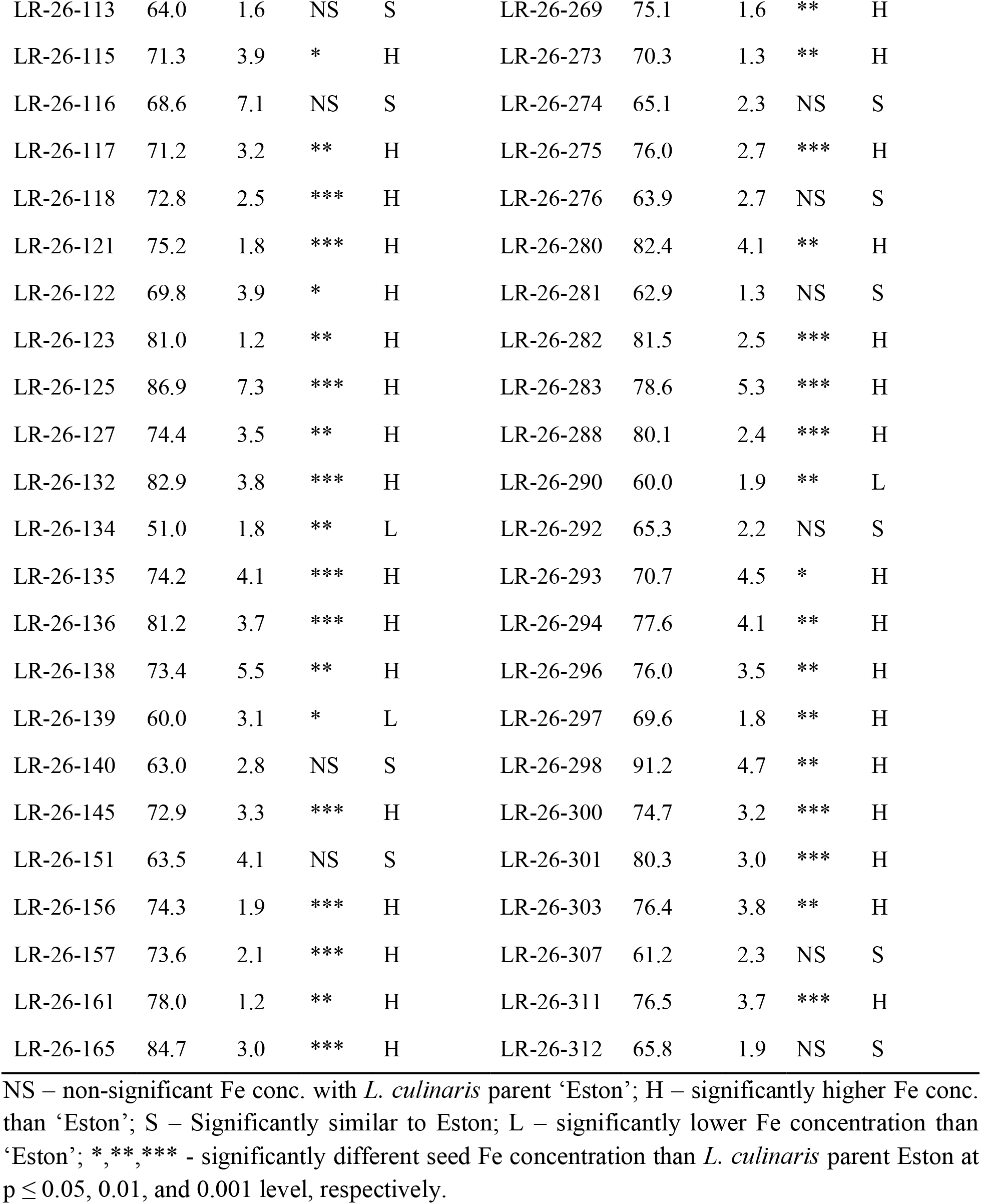
Seed Fe concentration of LR-26 interspecific RILs and their parents grown at Sutherland farm (2014 and 2015) and at Crop Science Field Laboratory, University of Saskatchewan in 2015.

